# Mitoguardin-2 is a lipid transporter and its lipid transfer ability is required for function

**DOI:** 10.1101/2022.07.08.499339

**Authors:** Zhouping Hong, Jyoti Adlakha, Emily Guinn, Fabian Giska, Kallol Gupta, Thomas J. Melia, Karin M. Reinisch

**Affiliations:** Department of Cell Biology, Yale University School of Medicine. New Haven, CT 06520; Nanobiology Institute, Yale University, West Haven, CT06516

## Abstract

Lipid transport proteins at membrane contact sites, where organelles are closely apposed, are critical in redistributing lipids from the endoplasmic reticulum (ER), where they are made, to other cellular membranes. Such protein mediated transfer is especially important for maintaining organelles disconnected from secretory pathways, like mitochondria. Here we identify mitoguardin-2, a mitochondrial protein at contacts with the ER and/or lipid droplets (LDs), as a lipid transporter. An X-ray structure shows that the C-terminal domain of mitoguardin-2 has a hydrophobic cavity that binds lipids. Mass spectrometry analysis reveals that both glycerophospholipids and free-fatty acids co-purify with mitoguardin-2 from cells, and that each mitoguardin-2 can accommodate up to two lipids. Mitoguardin-2 transfers glycerophospholipids between membranes in vitro, and this transport ability is required for roles both in mitochondrial and LD biology. While it is not established that protein-mediated transfer at contacts plays a role in LD metabolism, our findings raise the possibility that mitoguardin-2 functions in transporting fatty acids and glycerophospholipids at mitochondria-LD contacts.

## Introduction

Membrane contacts, sites where organelles are closely apposed, have emerged as a major means of intraorganellar communication and regulation (Lees et al., 2017; Prinz et al., 2020). These sites are important for membrane lipid homeostasis, including for lipid transfer between organelles, and especially to/from organelles disconnected from vesicle trafficking pathways, such as mitochondria. Proteins localized to contacts solubilize and transfer lipids across the cytosol between organellar membranes. Some such proteins mediate bulk lipid transfer, for example for membrane expansion, while others transport specific lipids, such as phosphoinositide lipids, to tweak the lipid compositions of existing membranes (Reinisch and Prinz, 2021). One approach to better understanding the physiological roles of contact sites and the processes that occur there is to characterize protein residents of these sites.

Here we characterize mitoguardin-2 (MIGA2), a mitochondrial protein present across most tissue types in higher eukaryotes, as a lipid transfer protein. MIGA2 localizes to contact sites between mitochondria and the ER or mitochondria and lipid droplets (LDs)(Freyre et al., 2019). It comprises an N-terminal transmembrane segment, which anchors it in the outer mitochondrial membrane, followed by a coiled-coil embedded in a largely unstructured linker, and a folded module at its C-terminus (Fig. 1A). An FFAT (two phenylalanines in an acidic tract) motif within the linker region of human MIGA2 binds the ER-proteins VAP-A/B to promote the formation of mitochondrial-ER contacts (Freyre et al., 2019). The C-terminal portion of MIGA2 is reported to promote contacts to LDs (Freyre et al., 2019). MIGA2 plays a role in mitochondrial health as its lack leads to mitochondrial fragmentation (Zhang et al., 2016), and in mammals it is implicated in lipid droplet biogenesis (Freyre et al., 2019). It is involved in the differentiation of and de novo lipogenesis in white adipocytes, promoting triglyceride synthesis and lipid droplet formation by still unclear mechanisms (Freyre et al., 2019). We show that a channel in the C-terminal domain of MIGA2, lined by hydrophobic residues, enables this domain to solubilize lipids and to transfer them between membranes in vitro, and that this ability is essential for MIGA2 functions in vivo, including in both mitochondrial and LD biology.

**Figure 1.**
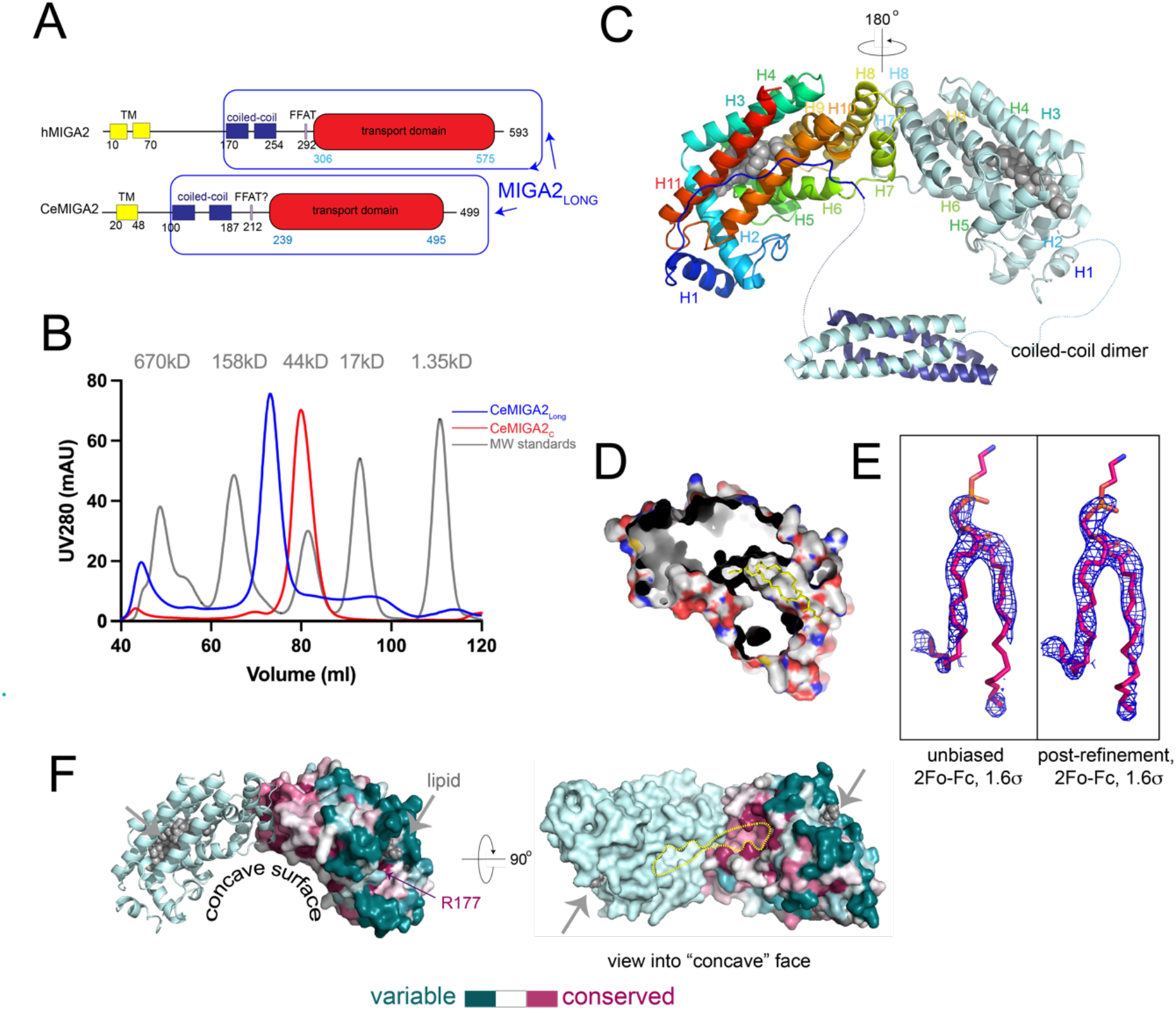
The structure of MIGA2. (A) Schematic of MIGA2 domain architecture. (B) Size exclusion profiles (Superdex 200 16/60) of CeMIGA2_long_ and CeMIGA2_C_, along with size standards (grey). CeMIGA2_long_ is a dimer, whereas CeMIGA2_C_ elutes more slowly as a monomer. hMIGA2 constructs behave the same (not shown).(C) Ribbons diagram of the CeMIGA2_long_ dimer that was crystallized, including the lipid transport module (colored from blue at the N-terminus to red at the C-terminus in one copy) and the 4-helix bundle that mediates dimerization. MIGA2_C_ dimerizes in the crystal as indicated. (D) MIGA2_C_ is cut away to reveal parts of the lipid binding tunnel. Residues at the MIGA2_C_ surface are colored red, blue, and white for oxygens, nitrogens, and carbons to illustrate that the tunnel is lined with hydrophobic residues. A yellow lipid is shown in the lipid binding site. (E) Initial electron density into which glycerophospholipid was modeled and the same density after refinement. Both densities are from 2Fo-Fc difference maps contoured at 1.6 σ. (F) Conservation analysis of MIGA2_C_ (calculated in Consurf(Ashkenazy et al., 2016)). A MIGA2 dimer is shown and conservation is indicated for one of the monomers. Bound lipids are indicated. A strictly conserved arginine residue near the lipid headgroup, R177 in CeMIGA2, is indicated. The molecule is rotated to view the concave conserved surface former by the dimer. The “empty” end of the tunnels, where we did not find lipid density, are outlined in yellow. Additional lipids may be bound there. The interface between MIGA2_C_ domains, formed by helices H7 and H8, is highly conserved.

## Results and Discussion

### Its structure suggests MIGA2 as a lipid transfer protein

To analyze MIGA2’s function biochemically, we over-expressed in bacteria and purified cytosolic fragments of human and *C. elegans* MIGA2 comprising portions of the linker region and the C-terminal domain (hMIGA2_long_, CeMIGA_long_, Fig. 1A) or only the C-terminal domain (hMIGA2_C_, CeMIGA_C_). The C-terminal domain migrated as a monomer as assessed by size exclusion chromatography, whereas the longer fragments that included a coiled-coil in the linker region dimerized (Fig. 1B). Its propensity to dimerize can explain previous observations that MIGA2 overexpression causes mitochondrial clustering in cells (Zhang et al 2016).

We crystallized CeMIGA_long_ (residues 106-496), including the coiled-coil segment in the linker region as well as the C-terminal domain, and determined its structure at 3.3 Å resolution. The crystals belonged to spacegroup P3_1_21. We used an AlphaFold2 generated model of CeMIGA2_C_ as a search model for phasing by molecular replacement, finding two dimers comprising four such modules in each asymmetric unit. The domains in the MIGA2_C_ dimer are related by a 180° rotation (Fig. 1C). Maps additionally showed density for two four-helix bundles, each corresponding to a dimer of the coiled-coils from the linker region (Fig. 1C). Notably, the fold predicted for the MIGA2 C-terminal domain was not previously reported in the Protein Data Bank but is consistent with experimental data as assessed by the success of the molecular replacement strategy.

The all-helical C-terminal domain of MIGA2 is globular, measuring ∼65 × 35 × 35 Å, and features an L-shaped channel (∼895 Å^3^ in volume as calculated in CASTp (Tian et al., 2018)). The channel is lined with hydrophobic residues and so is suitable for solubilizing lipids (Fig. 1C-D). As MIGA2 localizes to contact sites, like most lipid transporters, this finding suggested a function for MIGA2 as a lipid transfer protein. Poorly defined density, which was not modeled, occupies one end of the channel. Difference density reminiscent of a glycerophospholipid at the other end of the channel was modeled as such (Fig. 1E). The bound lipid co-purified with the protein from the *E. coli* expression host and presumably corresponds to a mixture of glycerophospholipids present in this bacterium (primarily PE and PG). There was no density for the lipid headgroup, as might be expected if lipids are bound non-specifically. Residues within the channel were conserved evolutionarily (as analyzed by Consurf (Ashkenazy et al., 2016)), supporting their functional importance. Surface residues near the lipid headgroup (or where the lipid headgroup is expected to be) are highly variable, except for R177 (Fig. 1F). R177 is strictly conserved across species and appears to be important for lipid binding (see analysis in “Its lipid transfer ability is essential for MIGA2 function in vivo”); likely through interaction with the headgroup of lipid ligands. The MIGA2_C_ domain, which was shown to be important for LD targeting (Freyre et al, 2019), dimerizes to form a concave surface patch that is conserved (Fig 1F). We speculate that the conserved patch could mediate interactions with LDs via LD -associated proteins rather than directly with lipids.

We analyzed hMIGA2_long_ (residues 170-575), similarly produced in bacteria, by native mass spectrometry, finding that up to four lipids are bound per hMIGA2_long_ dimer, with masses corresponding to glycerophospholipid species present in *E. coli* (primarily PE and PG) (Fig. 2A). While we can unambiguously observe one lipid per monomer of CeMIGA2 in the crystal structure (Fig. 1E), it is not clear where the other lipid binds. A plausible explanation is that it binds in the section of the hydrophobic channel in which we see poorly defined difference density (Fig. 1E).

**Figure 2.**
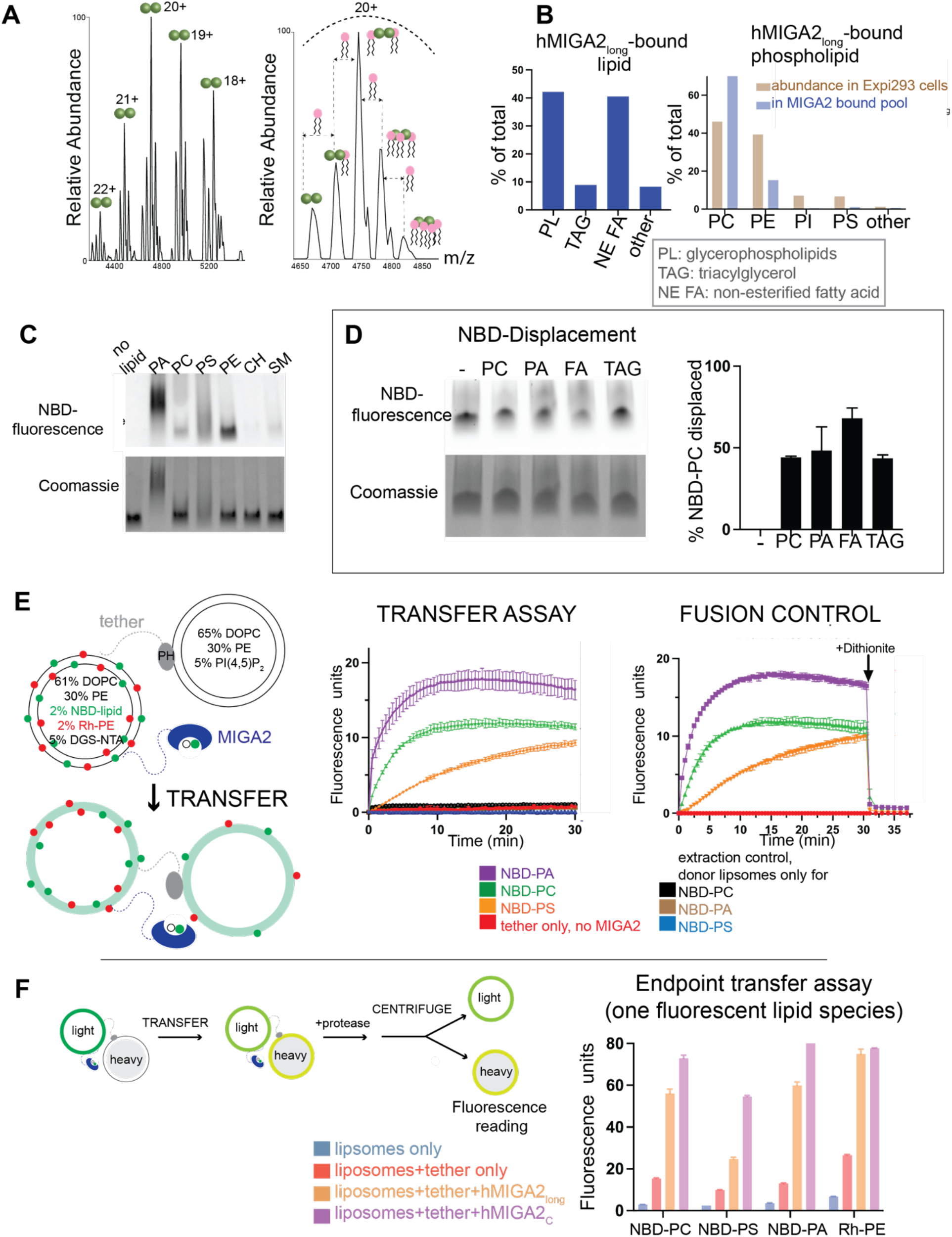
MIGA2 binds and transfers lipids between membranes. (A) Left: Native mass spectra of hMIGA2_long_-6x-his purified from bacteria, showing exclusive presence of the dimeric protein. Each charge state is further accompanied by a series of up to four lipid bound peaks. Right: Expansion of the 20+ charge state of native MS for hMIGA2_long_-6xhis dimer, showing up to four glycerophospholipids bound, with the average mass of 752 ± 5 Da. (B) Untargeted lipidomics of 3xFLAG-hMIGA2_long_, purified from mammalian (Expi293) cells. At left: distribution of lipid classes bound to hMIGA2_long_. At right: Distribution of glycerophospholipids bound to hMIGA2_long_ as compared to their abundance in cells (Lees et al., 2017). (C) 3xFLAG-hMIGA2_long_ was incubated with NBD-labeled lipids and examined by native PAGE. Phospholipids, visualized by their fluorescence, comigrated with protein, visualized by Coomassie blue staining. CH, cholesterol; SM, sphingomyelin. Because migration rates on native gels depend on the mass/charge ration of the sample, and because MIGA2 dimers can bind multiple lipids at once, MIGA2 incubated with charged lipids migrates as multiple species and is smeared. (D) hMIGA2_long_-6xhis was incubated with a 1:1 molar ratio of NBD-PC and unlabeled lipid and examined by native PAGE. (E) In the FRET-based transfer assay, donor and acceptor liposomes (compositions indicated) were tethered together in the presence or absence MIGA2 linked to the donor liposomes. The donor liposomes initially contain Rh-PE and NBD-lipids (PA, PC, PS), where FRET between the Rh and NBD initially reduces NBD fluorescence. As lipids are transferred from donor to acceptor liposomes, the Rh- and NBD-labeled lipids are diluted, resulting in reduced FRET and an increase in NBD-fluorescence. hMIGA2_long_ can transport NBD-PA, NBD-PC (acyl chain-labeled), and NBD-PS (headgroup-labeled). A donor only control shows that the fluorescence increase was due to transfer of lipids between liposomes rather than solely lipid extraction by hMIGA2. After the transfer reaction was completed, dithionite was added to rule out the possibility of fusion between donor and acceptor liposomes, which would also result in a fluorescence increase. The fluorescence reduction after dithionite addition is the same for reactions containing hMIGA2 as those without, indicating that fusion has not occurred. Each experiment was performed in triplicate. SDs are shown. (F) In the end-point transfer assay, “heavy” acceptor liposomes contain 0.75 M sucrose; donor liposomes have a single species of fluorescently labeled lipid. After transfer (30 min), proteinase K was added to digest proteins, the light donor and heavy acceptor liposomes were separated by centrifugation, and the fluorescence increase of the heavy acceptor liposomes monitored. Both hMIGA2_long_ and hMIGA2_C_ transport NBD-PC, -PS, -PA, and Rh-PE. Initial fluorescence readings before transport were the same for all samples. Each experiment was performed in triplicate. SDs are shown.

### MIGA2 binds lipids and can transfer them between membranes

As MIGA2 is present only in higher eukaryotes, which have different membrane compositions from bacteria, we further analyzed by LC/LC/MS lipids that co-purified with hMIGA2_long_ expressed and isolated from mammalian cells (Expi293F). The protein used in these experiments was purified using size exclusion chromatography to remove as much as possible non-specifically bound lipids (Maeda et al., 2013). We found that MIGA2 associated with glycerophospholipids (40 mole % of all lipids bound), fatty acids (40 mole % of all lipids bound), and triglycerides (10 mole % of all lipids bound). Of the glycerophospholipids, MIGA2 bound mostly PC and PE, with PC enriched slightly as compared to its abundance in cells (Fig. 2B). PS and PI represented a small fraction of the glycerolipids bound. The finding that MIGA2 associates with PC, FA and TAG is intriguing given the association of MIGA2 with LDs at contact sites, where LD’s comprise a neutral lipid core (TAG and cholesterol esters) surrounded by a monolayer of mostly PC. In cells, TAGs are hydrolyzed to FAs for transport to mitochondria for energy production or to the ER as precursors for glycerol- and glycerophospholipid synthesis. Thus, a possibility is that MIGA2 facilitates fatty acid transfer at mitochondria-LD contacts.

To further assess which lipids MIGA2 might bind, we incubated hMIGA2_long_ with NBD-labeled lipids, ran the samples on a native gel, and visualized bound lipids based on their NBD-fluorescence. NBD-labeled PC, PS, PE and PA are commercially available and co-migrated with hMIGA2_long_; we did not observe significant co-migration with NBD-labeled sterols or sphingomyelin, consistent with the mass spectroscopy analysis described above (Fig. 2C). The finding that MIGA2 binds PA is of interest as PA plays important roles in mitochondrial biology as a signaling molecule that regulates mitochondrial dynamics and as a precursor for cardiolipin. That PA was not detected among the lipids that co-purified with hMIGA2_long_ (as analyzed by lipidomics, above) reflects that it is present only in minute quantities in cells (van Meer and de Kroon, 2011). An important caveat in interpreting these experiments is that the NBD-modification used to visualize the lipids may affect their affinity for MIGA2. However, in a competition experiment, where hMIGA2_long_ was incubated with a 1:1 ratio of NBD-PC and an unmodified lipid, PA displaces half of the NBD-PC, confirming that MIGA2 binds unmodified PA robustly (Fig. 2D). Similarly, consistent with the lipidomics analysis, MIGA2 can bind FA and TAG as they also displace NBD-PC.

We next used a well established FRET-based assay (Kumar et al., 2018; Lees et al., 2017; Saheki et al., 2016) to follow whether MIGA2 can transfer lipids between membranes (Fig. 2E). To mimic contact sites, we tethered donor to acceptor liposomes using a previously described linker construct (Bian et al., 2018): it has an N-terminal hexahistidine tag that binds Ni-NTA lipids (5%) in the donor liposome and a C-terminal pleckstrin homolog (PH) domain that binds the phosphoinositide PI(4,5)P_2_ (5%) in the acceptor liposomes. Further, a hexahistidine tag was linked to the C-terminus of hMIGA2_long_ used in the transfer experiments, allowing robust association with the donor liposomes via the Ni-NTA lipids there. Initially, the donor liposomes contain Rh-PE (2%) and an NBD-labeled lipid (2%), whereas the acceptor liposomes lack fluorescently labeled lipids. FRET between lipid-associated Rh and NBD in the donor liposomes quenches NBD fluorescence. The addition of a lipid transfer protein and consequent transfer of one or both fluorescent lipid species to the acceptor liposomes results in their dilution, decreased FRET between Rh and NBD, and increased NBD fluorescence. As expected if MIGA2 can transfer fluorescently labeled lipid species, MIGA2 addition led to an increase in fluorescence. Lipid transfer was less efficient for NBD-PS than for NBD-PA or NBD-PC, which might be interpreted as indicating that MIGA2 has a preference for PA or PC over PS. An important caveat preventing this conclusion is that the assay assesses the transfer of NBD-modified rather than natural lipids. Further, the nature of the modification—whether the fluorescent group is conjugated to a fatty acyl moiety in the tail (NBD-PC, NBD-PA) or to the headgroup (NBD-PS, Rh-PE) – likely affects transport rates.

In a parallel study, we tethered “light” donor liposomes, which included a single Rh- or NBD-tagged fluorescent lipid species (2%), but not both at once, to “heavy” sucrose-filled acceptor liposomes initially lacking fluorescent lipids, and added MIGA2_long_ or hMIGA2_C_ to allow for transfer; then, to terminate any transfer reactions (after 30 minutes), we added protease (proteinase K) to degrade both the tether construct and MIGA2 (Fig. 2F). The heavy acceptor liposomes were separated from the light donor liposomes by centrifugation, and lipid transfer to the acceptor liposomes was assessed by their Rh- or NBD-fluorescence. Based on these experiments, MIGA2 (both hMIGA2_long_ and hMIGA2_C_) can bind and transport glycerophospholipids, including Rh-PE, NBD-PC, NBD-PA, and NBD-PS. As before, NBD-PS (headgroup modification) was transferred less efficiently than either NBD-PA or NBD-PC (acyl chain modification).

Thus, the contact site protein MIGA2 can bind glycerolipids and FAs within a hydrophobic channel and robustly transfers glycerolipids and possibly also FAs, although this remains to be tested, between membranes in vitro. As this manuscript was in preparation, Kim et al (Kim et al., 2022) reported the structure of MIGA2_C_ (from zebra fish) and used FRET-based assays similar to ours, with fluorescently modified lipids, to show that it transfers glycerophospholipids in vitro. They observed robust lipid transfer only in the case of NBD-PS (tail labeled), not NBD-PA, -PC, or -PE, and suggested that MIGA2 preferentially transfers PS. Their experimental setup differs most significantly from ours in that they did not use a tethering strategy for MIGA2 (there are other differences as well). Tethering MIGA2 to the liposomes assures that the protein interacts with the liposome, providing an opportunity for it to capture/extract lipid substrate, a stochastic event that determines the rate of the lipid transfer reaction. In the absence of a tether, whether a protein stays associated with membrane long enough for efficient lipid extraction depends heavily on the donor membrane composition (for example, see (Horenkamp et al., 2018; Zhang et al., 2022)); often the presence of acidic lipid (they used 10% NBD-PS, much more than would be present in ER, mitochondria, or LD’s) enhances protein interaction with membranes. We suspect that the transfer rates they observed reflect differences in lipid extraction rates arising from different donor liposome compositions and membrane characteristics, and that transfer of NBD-PC, -PA, and -PE would be enhanced upon addition of unlabeled PS to the donor liposomes. As noted, in our experimental setup both NBD-PC and NBD-PA transfer is robust. Further, arguing that PS is not a selective cargo, PS is not enriched in the pool of glycerolipids that co-purifies with MIGA2 in our lipidomics analysis. This is in contrast to other PS-specific transfer proteins like Osh6 (Maeda et al., 2013). Thus, whether MIGA2 preferentially transfers specific lipids remains an open question to be addressed in future through a combination of additional carefully controlled in vitro experiments and, equally important, studies in cells.

### Its lipid transfer ability is essential for MIGA2 function in vivo

Like many other lipid transport proteins, MIGA2 was first identified as a tether that promotes contact site formation. As a first step in determining whether, in addition to its tethering ability, MIGA2’s lipid transfer activity might be biologically important, we designed lipid transport incompetent mutants for use in functional studies. We introduced clusters of hydrophilic residues into the lipid binding channel of human MIGA2 so as to interfere with lipid solubilization (M1: V376D, L438E, F495H, I528K; M2: V381E, V502K, L506D, F524H) (Figure 3A). In another construct, we mutated the strictly conserved arginine near the lipid headgroup in the crystal structure (M3: R357N, S428G, where R357and S428 in hMIGA2 correspond to R177 and S248 in CeMIGA2, respectively). As assessed in the in vitro FRET-based assay described above with NBD-PS, all three mutant constructs (M1, M2, M3) are lipid transfer incompetent (Fig. 3B).

**Figure 3.**
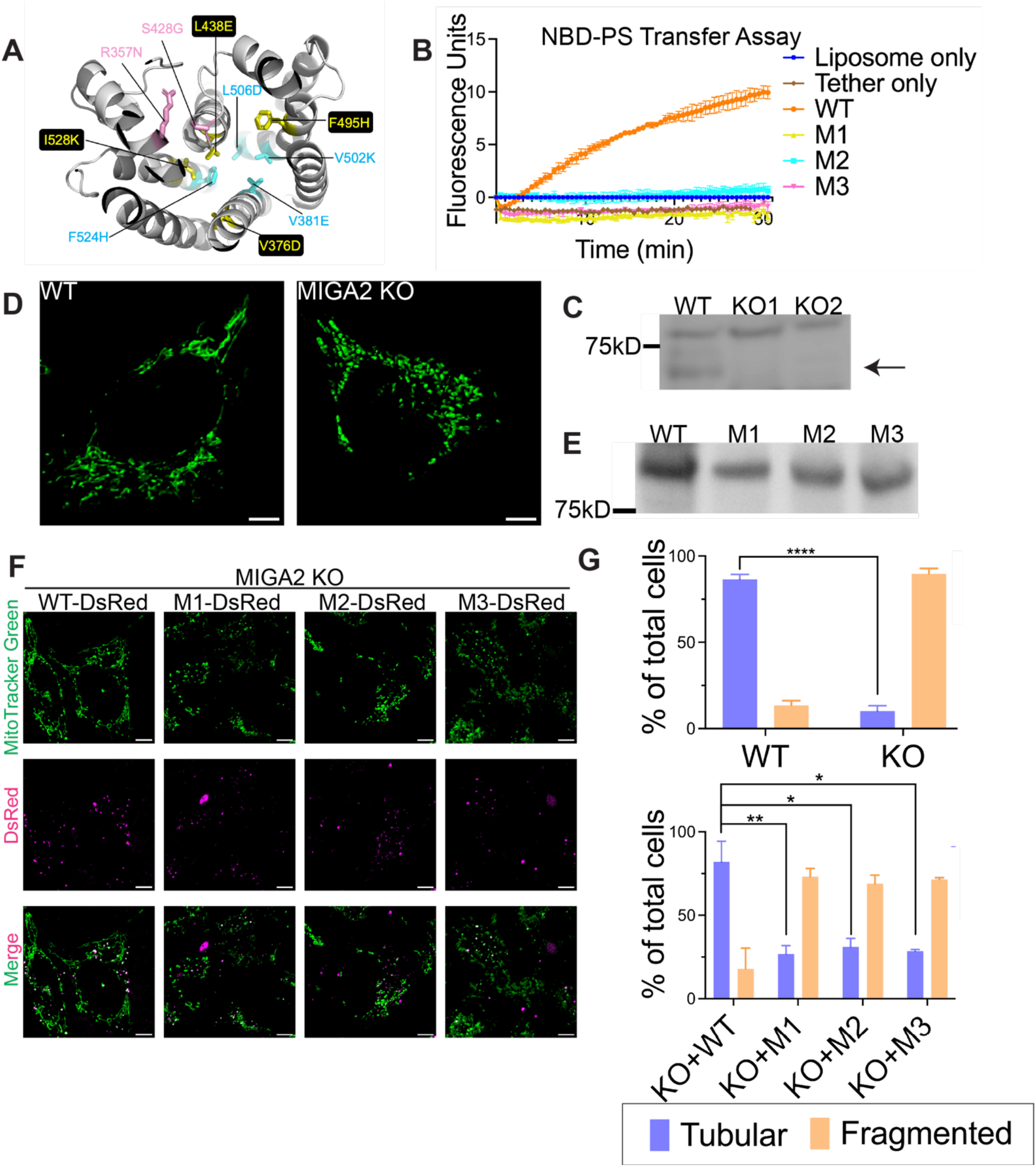
MIGA2’s lipid transfer ability is essential for its function in mitochondria. (A) In two mutant forms of hMIGA2 (M1 and M2), hydrophobic residues in the lipid binding channel were altered to hydrophilic amino acids, indicated in yellow and cyan, respectively. In a third mutant (M3), in pink, a strictly conserved residue near the lipid headgroup was changed to asparagine (R357N in hMIGA2). The hMIGA2_C_ shown was modeled in AlphaFold2. (B) All three mutants lose the ability to transfer lipids in the FRET-based assay with NBD-PS. Each experiment was performed in triplicate. SDs are shown. (C) Knockout of MIGA2 in Hela cells was confirmed by Western blot against MIGA2. The upper non-specific band served as an internal loading control. The arrow indicates the band of MIGA2. (D) Mitochondria stained with MitoTracker Green in WT Hela cells are tubular, while in MIGA2 KO cells are fragmented. Scale bar, 5μm. (E) Expression of WT MIGA2-DsRed and M1, M2, and M3-DsRed mutants in MIGA2 KO cells was compared by Western blot against DsRed. The four constructs expressed at similar level. (F) Expression of WT MIGA2-DsRed in the KO cells rescued the mitochondria morphology, whereas cells expressing either M1-, M2-, or M3-DsRed, the lipid transfer incompetent constructs, showed fragmented mitochondria. Scale bar, 5μm. (G) Quantification of tubular and fragmented mitochondria in different cells. Statistical significance was calculated by Welch’s two-tailed unpaired t test. Results were indicated in the following manner: * for p < 0.05, ** for p < 0.01, **** for p < 0.0001, where p < 0.05 is considered as significantly different. Graphs in the same panel are displayed with the same brightness and contrast settings.

Zhang et al (2016) showed that mitochondria are fragmented in MIGA2 knockout (KO) cells. As a tool to assess whether a lack of MIGA2 lipid transport ability might be responsible for this mitochondrial phenotype, we used CRISPR/Cas9 technology to make a MIGA2 KO Hela cell line (Fig. 3 C). We stained mitochondria with MitoTracker Green in WT cells, the KO cells, and cells transiently expressing WT MIGA2-DsRed or mutated versions (M1-DsRed, M2-DsRed,or M3-DsRed), then imaged them. 90% of MIGA2 KO cells had fragmented, dot-like mitochondria, significantly different from WT cells, which mostly had tubular mitochondria (∼90%) (Fig. 3 D, G). More importantly, transiently expressing WT MIGA2-DsRed in the KO cells rescued mitochondrial morphology in 80% of the cells, whereas only ∼30% of the cells expressing the lipid transfer incompetent constructs (M1-DsRed, M2-DsRed, or M3-DsRed) had tubular mitochondria (Fig. 3 F,G). To avoid mitochondria clustering caused by MIGA2 overexpression (Zhang et al., 2016), we used a relatively weak promotor, PGK to exogenously express MIGA2-DsRed. The expression levels for the transiently expressed MIGA2 WT and mutants were similar (Fig. 3 E). This indicates that the lipid transport ability of MIGA2 is necessary for its function in mitochondria.

Freyre et al (2019) discovered that MIGA2 targets to lipid droplets and participates in their formation. Like them, we found that depletion of MIGA2 in the KO cells reduced the number of lipid droplets and decreased the number of large lipid droplets after oleic acid (OA) treatment (Fig. 4A, C). The phenotype of fewer and smaller LDs in MIGA2 KO cells was reversed in cells expressing MIGA2-DsRed (Fig. 4 B, C). In contrast, expression of MIGA2 mutants deficient for lipid transport failed to increase the number and size of LDs in MIGA2 KO cells (Fig. 4 B, C), suggesting a role for MIGA2’s lipid transport ability in LD formation.

**Figure 4.**
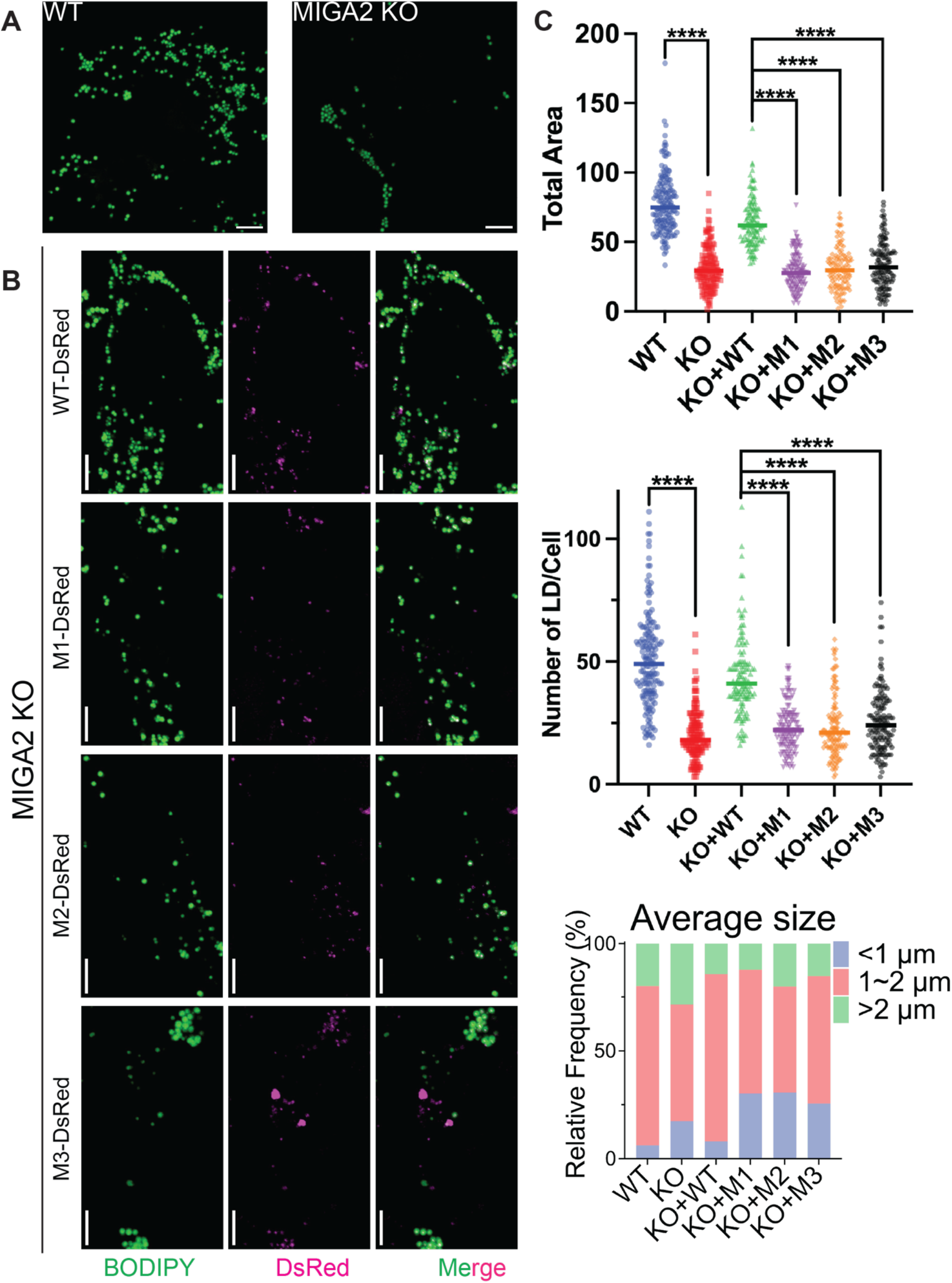
MIGA2’s lipid transfer ability is essential for its function in lipid droplet biology. (A) Lipid droplets were stained with BODIPY green 493/503 and imaged in cells treated with 100μM OA for 18-24 hours. MIGA2 KO Hela cells had fewer and more small lipid droplets than the WT cells. Scale bar, 5μm. (B) phenotype of fewer and smaller LDs in MIGA2 KO cells was reversed in those cells expressing MIGA2-DsRed. In contrast, MIGA2 mutants failed to do so. (C) Quantification of tubular and fragmented mitochondria in different cells. Statistical significance was calculated by Welch’s two-tailed unpaired t test. Results were indicated in the following manner: * for p < 0.05, ** for p < 0.01, **** for p < 0.0001, where p < 0.05 is considered as significantly different.

### Possible mechanisms for MIGA2 function

The lipid transport ability of MIGA2 is thus intrinsic to its functions both in mitochondrial and LD biology. Mitochondria rely on protein-mediated lipid transport at contact sites, especially with the ER, for most of their membrane lipids as they are disconnected from vesicle trafficking pathways. A number of lipid transport proteins have already been identified at ER-mitochondrial contacts, and it will be important to discover a distinguishing role for MIGA2 at these sites. Perhaps it supplies mitochondria with PA, or perhaps as proposed by Kim et al it participates in PS transfer to the mitochondria, where PS is a precursor for PE; or maybe it can transfer PE from mitochondria back to the ER. Or MIGA2 could act as a non-specific glycerophospholipid transporter that equilibrates the lipid compositions of ER and mitochondrial membranes at contact sites. LD-mitochondrial contacts have been implicated in LD biogenesis previously (Benador et al., 2018). That lipid exchange takes place at these sites is not well established (but see (Du et al., 2020)). It is nevertheless possible that MIGA2-mediated lipid transport there plays a role in LD formation; or maybe MIGA2 acts at three-way contacts between the LDs, mitochondria, and the ER, where de novo LD formation takes place (reviewed in (Choudhary and Schneiter, 2021; Henne et al., 2020; Olzmann and Carvalho, 2019)). To us, a very interesting possibility, given that MIGA2 co-purifies with, binds and transfers PC (the prominent glycerophospholipid in the monolayer surrounding the LD core), and binds FA’s (produced from TAGs in the LD core), is that it may facilitate PC and FA transport between LDs and mitochondria at LD-mitochondrial contacts and FA transport on to other organelles from there, to play a role in ATP generation (mitochondria) or else as a precursor for glycerol- and glycerophospholipid synthesis (ER). To our knowledge, no FA transfer protein has yet been identified. Further studies that identify the physiological ligands of MIGA2 are the next step in understanding the molecular basis of MIGA2 function.

## Materials and Methods

### Materials

Rabbit polyclonal anti MIGA2 antibody (ab 122713) and rabbit polyclonal anti DsRed antibody (ab167453) were purchased from abcam. Almost all lipids were purchased from Avanti Polar Lipids: DOPC (850375), liver PE (840026), DGS-NTA (Ni) (709404), Liss Rhod PE (810150), Brain PI (4,5) P2 (840046), NBD-PA (810198), NBD-PS (810198), NBD-PC (810133), NBD-PE (81015),NBD-cholesterol (810250), NBD-sphingomylein (810219), 16:0 PA (830855), triolein (18:1 TG) (870110). Palmitic acid (P0500) was ordered from Sigma Aldrich. Sodium dithionite was from Sigma Aldrich (157953). The Hela cell line was a gift from Mals Mariappan (Yale University, New Haven, CT), *C. elegans* cDNA was a gift from Daniel Colon-Ramos (Yale University, New Haven, CT). Plasmid for the 6xhis-PH-tethering construct (Bian et al., 2018)was a gift from Pietro De Camilli (Yale University, New Haven, CT).

### Plasmid construction

*C*.*elegans* MIGA (Uniprot Q21096) fragments were amplified from *C*.*elegans* cDNA library and subcloned into pET28-6xhis-sumo plasmid or pET29 or pCMV10 plasmid with C-terminal 6xhis tag or N-terminal 3xFLAG tag. Human MIGA2 (Uniprot Q7L4E1) gene was codon-optimized for bacterial expression and synthesized by Genescript. The gene fragments (170-593, 306-593) were PCR amplified and cloned into pCMV10 or pET29 plasmid with N-terminal 3xFLAG or a C-terminal 6xhis tag. The mutant gene fragments were synthesized by Genescript and PCR amplified and cloned into either pCMV10 plasmid with a N-terminal 3xFLAG and a C-terminal 6xhis tag for biochemical experiments or pLENTi-PGK plasmid with a C-terminal DsRed tag for imaging.

### Protein expression and purification

#### For crystallization

The pET28-6xhis-sumo-CeMIGA(106-496) construct was expressed in C43 (DE3) *Escherichia coli* cells. Cells were grown at 37°C to an OD_600_ of 0.6–0.8, when protein expression was induced with 0.8 mM IPTG, and then cells were cultured at 18°C for another 6 h. Cells were pelleted, resuspended in buffer A (20 mM HEPES, pH 7.4, 200 mM NaCl, 1mM TCEP, and 5% glycerol) containing 1× complete EDTA-free protease inhibitor cocktail (Roche 1187358001) and lysed in an Emulsiflex-C5 cell disruptor (Avestin). Cell lysates were clarified via centrifugation at 27,000 g for 30 min. To collect the protein, supernatant was incubated with Ni-NTA resin (QIAGEN 30210) for 1 hour at 4°C, and then the resin was washed tandemly with buffer B (buffer A + 0.005% Triton X-100), buffer C (buffer B + 10mM imidazole), buffer D (buffer B + 20mM imidazole), buffer E (buffer B + 40mM imidazole), buffer A to remove extra Triton X-100. Retained protein was eluted from the resin with buffer A supplemented with 250 mM imidazole. Sumo protease was added to digest overnight at 4°C. After removing imidazole by buffer exchange, the sample was bound to the Ni-NTA resin again to remove protease and 6xhis-sumo. The flow-through was collected, concentrated in a 30-kD molecular weight cutoff (MWCO) Amicon centrifugal filtration device, and loaded onto a Superdex 200 16/60 column (GE Healthcare) equilibrated with buffer F (20 mM HEPES, pH 7.4, 200 mM NaCl, 1mM TCEP). Peak fractions containing pure MIGA were recovered and concentrated.

#### For biochemical assays

3xFLAG-MIGA2_long_ WT and mutants for Expi293 cells expression were transfected into Expi293F cells according to manufacturer instructions for 48 hours. Cells were pelleted and resuspended in buffer A and lysed by 5 freeze-thaw cycles. Cell lysates were clarified via centrifugation at 27,000 g for 30 min, and the supernatant was incubated with preequilibrated anti-FLAG M2 affinity resin (Sigma-Aldrich, A2220) for 2 hours at 4°C. The resin was washed with buffer A and incubated overnight with buffer A containing 2.5 mM ATP and 5 mM MgCl_2_. The protein was eluted with buffer A supplemented with 0.2 mg/ml 3× FLAG peptide (APExBio A6001), concentrated in a 30-kD MWCO Amicon centrifugal filtration device, and loaded onto a Superdex 200 10/30 column (GE Healthcare) equilibrated with buffer A. Peak fractions were pooled and concentrated.

MIGA2_long_-6xhis and MIGA2_C_-6xhis were expressed in BL21 (DE3) *Escherichia coli* cells. The expression and purification of these constructs were the same as that for CeMIGA(106-496) except that no detergent added during washes, and no sumo digestion and rebinding, instead loading onto Superdex 200 16/60 column directly after elution and concentration.

The 6xhis-PH-tether construct was purified as described before (Bian et al., 2018).

### Protein crystallization, structure determination, and refinement

Crystals of *C. elegans* MIGA (106-496) at 6 mg/ml were grown at 18°C using the sitting-drop vapor-diffusion method. Equal volumes of protein and reservoir solution (0.2 M sodium malonate, pH 7.4, 22% PEG335) were mixed. Crystals, which belonged to spacegroup P3_1_21 (a=91.3, b=91.3, c=366.74 Å), were transferred to solutions that also included cryo-protectants, and flash frozen in liquid nitrogen. We examined ∼100 crystals. Although most did not diffract to atomic resolution, a small number diffracted to ∼4 Å or better. We found a single crystal among them that were cryo-protected in 25% ethylene glycol that diffracted to 3.3 Å, from which we collected the data set used in structure determination. Diffraction data were collected at the Northeastern Collaborative Access Team (NE-CAT) beamline 24-ID-C at the Advanced Photon Source, using a Dectris EIGER2 × 16M pixel array detector, and processed using XDS (Kabsch, 2010). Statistics for data collection are shown in Table 1. For phasing, we used molecular replacement with Phaser MR (McCoy et al., 2007) using a CeMIGA2_C_ model generated by AlphaFold2 Colab (Jumper et al., 2021) preprocessed with phenix.process_predicted_model (Liebschner et al., 2019) to remove the low confidence loop regions. Four copies of CeMIGA2_C_ were placed in the asymmetric unit. Initial maps showed clear density for two four-helix bundles, representing dimers of the coiled-coil helices from the CeMIGA2 linker. The helices were modeled manually in Coot (Emsley et al., 2010). Additional unmodeled density was observed in the Fo-Fc maps within the hydrophobic cavity in two MIGA2_C_ domains, which clearly corresponded to bound glycerophospholipids. We built PE 16:0 (chemical id PEF) into these densities and eventually, using non-crystallographic symmetry, placed PE into less well defined densities in the remaining two monomers. The refinement consisted of cycles of manual rebuilding in Coot and automated refinement in Phenix(Liebschner et al., 2019), including isotropic b-factors, Translation-Libration-Screw refinement and non-crystallographic symmetry restraints. Refinement statistics are in Table 1. The coordinates and structure factors will be deposited in the Protein Data Bank (accession no. ####).

**Table 1:** Data collection and refinement statistics

|  |  |
| --- | --- |
|  | CeMIGA2<sub>long</sub> |
| Wavelength (Å) | 0.979110 |
| Space group | P 3<sub>1</sub> 2 1 |
| *Cell dimensions* |  |
| a, b, c (Å) | 91.3, 91.3, 366.7 |
| $\alpha$, $\beta$, $\gamma$ (°) | 90, 90, 120 |
| Monomers/ASU | 4 |
| *Data collection* |  |
| Resolution range (Å) | 48.36-3.30 (3.42-3.30) |
| Completeness (%) | 99.9 (99.2) |
| Redundancy | 10.2 (10.5) |
| I/$\sigma$(I) | 14 (1.6) |
| Rmerge (%) | 16.9 (73.6) |
| Rmeas (%) | 17.9 (77.4) |
| Rpim (%) | 5.6 (23.6) |
| CC (1/2) | 99.3 (83.5) |
| *Refinement* |  |
| Rwork (%) | 26.76 (32.82) |
| Rfree (%) | 29.41 (32.49) |
| RMSD bond angles/lengths | 0.642/0.005 |
| Ramachandran statistics (%) | 94.85/4.22/0.92 |
| PDB accession code | XXXX |

Values in parentheses correspond to the highest resolution shell. The Ramachandran statistics show the percentage of residues in favoured/allowed/other regions.

Figures were made using Pymol (The PyMOL Molecular Graphics System, Version 1.5.0.4. Schrödinger, LLC, New York.).

### Lipid Co-migration assay

Purified hMIGA2_long_ was mixed with 1μl of NBD-labeled lipids (at 1mg/ml) in 20 μl total reaction volumes and incubated on ice for 2 hours. Samples were loaded onto 4–15% Mini-Protean Precast Native gels and run for 90 minutes at 100 V. NBD fluorescence was visualized using an ImageQuant LAS4000 (GE Healthcare). Then gels were stained with Coomassie blue G250 to visualize total protein. Images were analyzed via Fiji (ImageJ).

### Lipid Competition assay

NBD–PC and another non-labelled lipids at a 1:1 molar ratio (1.13 mM each) were mixed with hMIGA2_long_ in 40 μl reaction volumes and incubated on ice for 2 hours. Samples were loaded onto 4–15% Mini-Protean Precast Native gels and run for 90 minutes at 100 V. NBD fluorescence was visualized using an ImageQuant LAS4000 (GE Healthcare).

### Liposome preparation

#### For FRET based assay

Lipids in chloroform were mixed (donor liposomes: 61% DOPC, 30% liver PE, 2% NBD-PA, 2% Rh-PE, and 5% DGS-NTA (Ni); acceptor liposomes: 65% DOPC, 30% liver PE, and 5% PI(4,5)P_2_) and dried to thin films and vacuumed for 30 minutes. Lipids were subsequently dissolved in buffer F at a total lipid concentration of 1 mM and incubated at 37°C for 1 hour, vortexing every 10-15 min. Liposomes were subjected to 10 freeze–thaw cycles alternating between liquid nitrogen and room temperature water bath with vortexing every three cycles. Crude liposomes were then extruded through a polycarbonate filter with 100 nm pore size a total of 11 times via a mini extruder (Avanti Polar Lipids) and used within 24 hours.

#### For End-point transfer assay

Light donor liposomes (63% DOPC, 30% liver PE, 2% fluorescently labeled lipids, and 5% DGS-NTA (Ni)) were prepared similarly as before except that either NBD-lipids or Rhodamine-PE, but not both, were included. For heavy acceptor liposomes, lipids in chloroform were mixed, dried and vacuumed as above. The lipid film was dissolved in buffer F containing 0.75 M sucrose. at a total lipid concentration of 1 mM and incubated at 37°C for 1 hour, vortexing every 10-15 min. Liposomes were subjected to 10 freeze–thaw cycles, and then mixed with 2 volumes of buffer F and pelleted for 40 min at 18,500 x g. The resulting liposome pellet was resuspended in buffer F and pelleted again for 30 min. The second pellet was resuspended to 1 mM total lipid concentration in buffer F and used immediately.

### In Vitro Lipid transfer assay

#### FRET based assay

Lipid-transfer experiments were set up at 30°C in 96-well plates, with 100-μl reaction volumes containing 200 μM lipids in donor liposomes and 200 μM lipids in acceptor liposomes. Proteins (0.25 μM each of tether and lipid transport protein) were added to start the reaction, and after excitation at 460 nm, NBD emission (538 nm) was monitored for 30 minutes using a Synergy H1 Multi-Mode Microplate Reader (Agilent).

#### Dithionite Assay

Lipid transfer assays were performed as above, except for the addition of freshly prepared dithionite (to 5 mM final concentration) after the last time point, and NBD fluorescence was monitored for an additional 5 min.

#### End-point transfer assay

Lipid transfer reactions were performed in 100 μl volumes. The reaction was initiated by adding 200 μM lipids each in light donor and in heavy acceptor liposomes into 0.25 μM protein (tether and hMIGA2_long_ or hMIGA2_C_) and terminated by the addition of 1 mg/ml proteinase K. The complete digestion of the proteins was confirmed by SDS-PAGE gel. Light and heavy liposomes were then separated by centrifugation at 18,500 g for 15 min and heavy liposomes recovered in the pellets were resuspended in 100 μl buffer F. Fluorescence signal (NBD: ex 440 nm, em 514 nm; Rhodamine: ex 550 nm, em 590 nm) present in the pellet was determined using a Synergy H1 Multi-Mode Microplate Reader (Agilent).

### Mass Spectrometry analysis of hMIGA2_long_

#### Lipidomic analysis

Gel filtrated 3xFLAG-hMIGA2_long_ was sent to Michigan State University’s Collaborative Mass spectrometry Core for untargeted lipidomics analysis. The sample was spiked with internal standards and calibration mixture. Lipids were extracted with MTBE twice, and after drying, resuspended in isopropanol containing 0.01% BHT. The sample was resolved by a Shimadzu Prominance HPLC and lipid species were detected by a Thermo Scientific LTQ-Orbitrap Velos mass spectrometer in both positive and negative ionization modes. Lipid species were quantified based on internal standards and summed by lipid class.

#### Native Mass spectrometry

The hMIGA2 sample was buffer exchanged to 200 mM ammonium acetate (MP Biomedicals), 2 mM DTT with Zeba Spin Desalting Columns (Thermo-Fisher Scientific). The protein concentration in the analyzed sample was in the range between 1 μM and 5 μM. Native-MS was performed on Q Exactive UHMR (Thermo-Fisher Scientific) using in-house nano-emitter capillaries. The tips in the capillaries were formed by pulling borosilicate glass capillaries (OD: 1.2 mm, ID: 0.69 mm, length: 10 cm, Sutter Instruments) using a Flaming/Brown micropipette puller (ModelP-1000, Sutter Instruments). Then the nano-emitters were coated with gold using rotary pumped coater Q150R Plus (Quorum Technologies). To perform the measurement the emitter filled with the sample was installed into Nanospray Flex Ion Source (Thermo-Fisher Scientific). MS parameters for the analysis of the proteins or protein complexes include spray voltage 1.1–1.3 kV, capillary temperature 275°C, resolving power 6250 at m/z of 400, ultrahigh vacuum pressure 4.6e-10–8.18e-10, in-source trapping between −100 V and -200 V.

### Functional Experiments

#### Cell culture and transfection

Hela cells were cultured in DMEM (Thermo Fisher Scientific 11965092) supplemented with 10% FBS (Thermo Fisher Scientific 10438062) and 1% penicillin-streptomycin (Thermo Fisher Scientific 15140122) at 37°C in a 5% CO2 incubator. Cells were used in experiments before passage 5. DNA transfection was performed using Lipofectamine 3000 (Thermo Fisher Scientific, L3000015) according to the manufacturer’s instruction.

#### Generation of MIGA2 KO cell line

CRISPR guide RNAs (gRNAs) targeting the third exons of MIGA2 were designed using the online TKO CRISPR Design Tool (https://crispr.ccbr.utoronto.ca/crisprdb/public/library/TKOv3/). 5’-GCGGAAAGTCCTCTTTGCCA-3’ was chosen as gRNA to knock out MIGA2. The gRNA was cloned into pX458 (Addgene 48138) as described previously (Ran et al., 2013). Hela cells were transiently transfected with the constructs containing gRNAs. After 48 h, the GFP positive individual cells were selected by flow cytometry (BD FACSAria) and seeded in 96-well plates for single clones. The clones were validated by genotyping and Western blotting. For genotyping, briefly, genomic DNAs from single clones were extracted using QuickExtract DNA extraction solution (Lucigen QE0905T), and PCR products containing the site of Cas9 cleavage site were generated using the following primers: 5’-TAGACCTCACCTTCTCGGCACT-3’; 5’- CCAATATCCCCAAGTAGAGAGTG-3’. PCR products were sequenced.

#### Western blotting

The samples were denatured by the addition of SDS sample buffer (125 mM Tris-HCl, pH 6.8, 16.7% glycerol, 3% SDS, 0.042% bromophenol blue) and heated at 95C for 5 min. The proteins were separated using denaturing SDS-PAGE and analyzed by immunoblotting. The proteins were transferred onto a polyvinylidene fluoride (PVDF) membrane. After the transfer the membranes were incubated in 5% nonfat dry-milk–TBST (20 mM Tris-HCl pH 7.6; 150 mM NaCl; 0.1% Tween 20) for 1 h at room temperature. The membranes were then washed three times and incubated with primary antibodies at 1:1000 dilution in 5% BSA-TBST over night at 4°C or at room temperature for 90 minutes. The membranes were washed three times and incubated for 60 min with secondary antibodies coupled to horse-radish peroxidase in TBST containing 5% nonfat dry-milk. The proteins of interest were developed by enhanced chemiluminescence (ECL, BioRad) and visualized using ImageQuant LAS4000 (GE Healthcare).

#### Live cell imaging

Cells were plated on glass-bottomed 35mm live cell imaging plate and transfected with desired plasmids the next day. After 1 day, cells were treated with 100 μM OA (CAYMAN, 29557) or same amount of BSA control (CAYMAN, 29556) for 18∼24 hours if imaging lipid droplets. 48 hours after transfection, Cells were incubated with BODIPY 493/503 (Thermo Fisher Scientific #D3922) at 10 *μ*g/ml or MitoTracker Green (Thermo Fisher Scientific M7514) at 75-100 nM for 30 minutes, washed with PBS, and subjected to further live-cell imaging in 1x live cell imaging solution (Thermo Fisher Scientific, A14291DJ).

At the Center for Cellular and Molecular Imaging Facility at Yale, imaging was performed with a 63× oil-immersion objective on an inverted Zeiss LSM 880 laser scanning confocal microscope with AiryScan, using Zen Black acquisition software. It has a 63× oil-immersion objective lens with NA = 1.4.. Z-stack images were taken. Images were acquired using Nikon Elements and analyzed in Fiji (ImageJ).

For high throughput imaging of lipid droplets, confocal microscopy was performed on Nikon Ti2-E inverted microscope with a 63x oil-immersion objective, CSUX1 camera Photometrics Prime 95B, Agilent laser 488 and 561 nm, and a temperature setting at 37 C by the Oko Lab control system. Images were acquired using Nikon Elements and analyzed in Fiji (ImageJ).

Different samples were imaged in one session with the same settings. All images were analyzed in ImageJ. All data categorizing mitochondrial morphology were scored blindly. For lipid droplets analysis, The Z-stack was maximum projected and the images were set at the same threshold. Lipid droplets were analyzed after converting to binary.

#### Statistical analysis

Statistical analysis between groups was performed using Prism (GraphPad Software) with Welch’s two-tailed unpaired t test. Results were indicated in the following manner: * for p < 0.05, ** for p < 0.01, **** for p < 0.0001, where p < 0.05 is considered as significantly different.

## Acknowledgments

We thank the staff of the NE-CAT beamlines at the Advanced Photon Source (Argonne National Laboratory) for their help in Data Collection. NE-CAT is funded by the National Institute of General Medical Sciences of the National Institutes of Health (NIH; P30 GM124165), NIH-ORIP HEI grant (S10OD021527), and contract DE-AC02-06CH11357. Our research was funded by grants from the NIGMS to KMR, TJM, and KG (R35GM131715, R01GM135290, R01GM141192) and support from the China Scholarship Council to ZH.

## Author contributions

ZH and KMR conceived the project; ZH designed and carried out all the experiments. JA helped with the structure determination and particularly model building and refinement; EG helped in developing imaging strategies. Native mass spectroscopy analysis was carried out by KG based on the spectrum generated by FG. KMR supervised all aspects of this work with help from TJM for functional assays, and wrote the manuscript with ZH, with inputs from TJM and KG and JA.

